# Development and In Vivo Evaluation of a Novel Bioabsorbable Polylactic Acid Middle Ear Ventilation Tube

**DOI:** 10.1101/2025.06.25.661110

**Authors:** Ying-Chang Lu, Chi-Chieh Chang, Ping-Tun Teng, Chien-Hsing Wu, Hsuan-Hsuan Wu, Chiung-Ju Lin, Yen-Hui Chan, Chen-Chi Wu

## Abstract

**Background:** Otitis media with effusion (OME) is a widespread condition causing hearing impairment, particularly in pediatric populations. Existing non-absorbable middle ear ventilation tubes frequently necessitate secondary surgical removal. Bioabsorbable polylactic acid (PLA) offers a promising alternative due to its inherent biocompatibility and tunable degradation characteristics. This study aimed to design, fabricate, and comprehensively evaluate a novel PLA middle ear ventilation tube.

**Methods:** Bioabsorbable PLA tubes were designed and fabricated based on commercial models. In vitro biocompatibility was assessed according to ISO 10993 guidelines. In vivo evaluations were performed in a guinea pig model, including otoscopic examinations, auditory brainstem response (ABR) measurements, micro-computed tomography (micro-CT) imaging, and histological analyses.

**Results:** The PLA tubes were successfully designed and fabricated, exhibiting dimensions comparable to commercially available products. In vitro testing confirmed their biocompatibility. In vivo observations demonstrated that the PLA segments remained stable, with no significant inflammation detected. ABR measurements revealed no adverse impact on hearing function. Micro-CT imaging confirmed tube integrity and displayed initial signs of degradation over a 9-month period. Histological analyses indicated a favorable tissue response with minimal foreign body reaction.

**Conclusion:** The developed PLA middle ear ventilation tube represents a highly promising alternative to conventional non-absorbable tubes. It demonstrates excellent biocompatibility, preserves auditory function, and exhibits a controlled degradation profile. This preclinical study provides strong support for further investigation and subsequent clinical trials to validate its safety and efficacy in human patients.

## 1. Introduction

Otitis media with effusion (OME) is a prevalent inflammatory condition characterized by the accumulation of fluid in the middle ear space without acute signs of infection. OME is a leading cause of hearing impairment in children, with an estimated prevalence of up to 10% in pediatric populations [1, 2]. The Eustachian tube, connecting the middle ear to the nasopharynx, is crucial for maintaining middle ear pressure equilibrium and drainage. Dysfunction of this tube, often due to immature development in children or conditions such as allergic rhinitis, sinusitis, or nasopharyngeal carcinoma in adults, can lead to OME [2]. Prolonged untreated OME (e.g., 2-3 months) can result in significant and potentially irreversible hearing loss, impacting speech development, learning, and overall quality of life [1].

For OME that does not respond to medical treatment, middle ear ventilation tubes are a common and effective surgical intervention [1, 3]. The procedure involves a small incision in the tympanic membrane (myringotomy) to drain fluid, followed by tube placement to maintain aeration and equalize pressure [4]. Unlike simple myringotomies, which heal rapidly, the tube prevents premature closure. This ensures prolonged ventilation (ideally 6–24 months) and fluid drainage while Eustachian tube function recovers [1, 4]. Over one million tympanostomy tube insertions are performed annually in the U.S., making it a leading pediatric surgical procedure [5]. The global clinical demand for these tubes highlights a critical need for improved solutions.

Current non-absorbable middle ear ventilation tubes, typically made from materials such as Teflon or silicone, have significant limitations [6]. A major concern is their failure to spontaneously extrude, reported in up to 40% of cases [7, 8]. Retained tubes can cause foreign body reactions, leading to granulation tissue, inflammation, and tympanosclerosis [9]. These complications often necessitate a second surgical procedure for tube removal, increasing healthcare costs and patient risks, especially for children who require general anesthesia [9]. These complications often necessitate a second surgical procedure for tube removal, increasing healthcare costs and patient risks, especially for children who require general anesthesia [10]. Patients with Eustachian tube dysfunction may develop retraction pockets from atrophic membranes, which are a precursor to cholesteatoma; perforations often require surgical repair. Rare but serious complications, such as opportunistic mycobacterial infections, have also been reported [11, 12]. These issues underscore the urgent clinical need for a novel middle ear ventilation tube that degrades predictably, thus eliminating the need for a second intervention.

In an effort to overcome the limitations of non-absorbable tubes, researchers have explored bioabsorbable alternatives. Previous attempts using materials such as poly-bis(ethylalanate)phosphazene, hyaluronic acid, and gelatin have shown promise in terms of biocompatibility and ventilation function. However, these materials often exhibited overly rapid absorption rates or caused inflammation, which limited their clinical applicability [13, 14]. Polylactic acid (PLA) is a promising bioabsorbable material for medical implants. It is FDA-approved, biocompatible, biodegradable, and derived from renewable plant sources [15]. Its degradation products—lactic acid, water, and carbon dioxide—are naturally metabolized and safely cleared, ensuring human safety [15, 16]. PLA’s widespread use in surgical sutures, bone fixation, and drug delivery demonstrates its reliability. Key advantages of PLA for middle ear ventilation tubes include its controllable degradation profile, which can be adjusted to precisely match the ideal one-to two-year ventilation period [3]. This eliminates the need for removal surgeries and minimizes long-term complications. PLA also possesses established biocompatibility (ISO 10993 certified). Its mildly acidic degradation environment may offer bacteriostatic properties, and it can be engineered with antimicrobial coatings or for drug loading. Given the advantages of PLA and the significant unmet clinical need, the objective of this study is to design, manufacture, and evaluate a novel bioabsorbable PLA middle ear ventilation tube. Our objective is to develop an optimized PLA tube with controlled degradation kinetics to effectively ventilate the middle ear.

## 2. Methods

### 2.1 Design and Fabrication of PLA Middle Ear Ventilation Tubes

The middle ear ventilation tube was designed to replicate the form and function of existing commercial products. As depicted in Figure 1, the tube features a dumbbell shape with a central lumen to facilitate exudate flow and pressure equalization. PLA was selected as the raw material due to its biodegradable and thermoplastic properties. A medical-grade stainless steel mold was developed to facilitate the injection molding process, utilizing a Sumitomo injection molding machine (se100d, Sumitomo, Japan). Injection molding was chosen for its suitability for mass production and its ability to ensure consistent product quality, both crucial for future product certification and commercialization.

**Figure 1.**
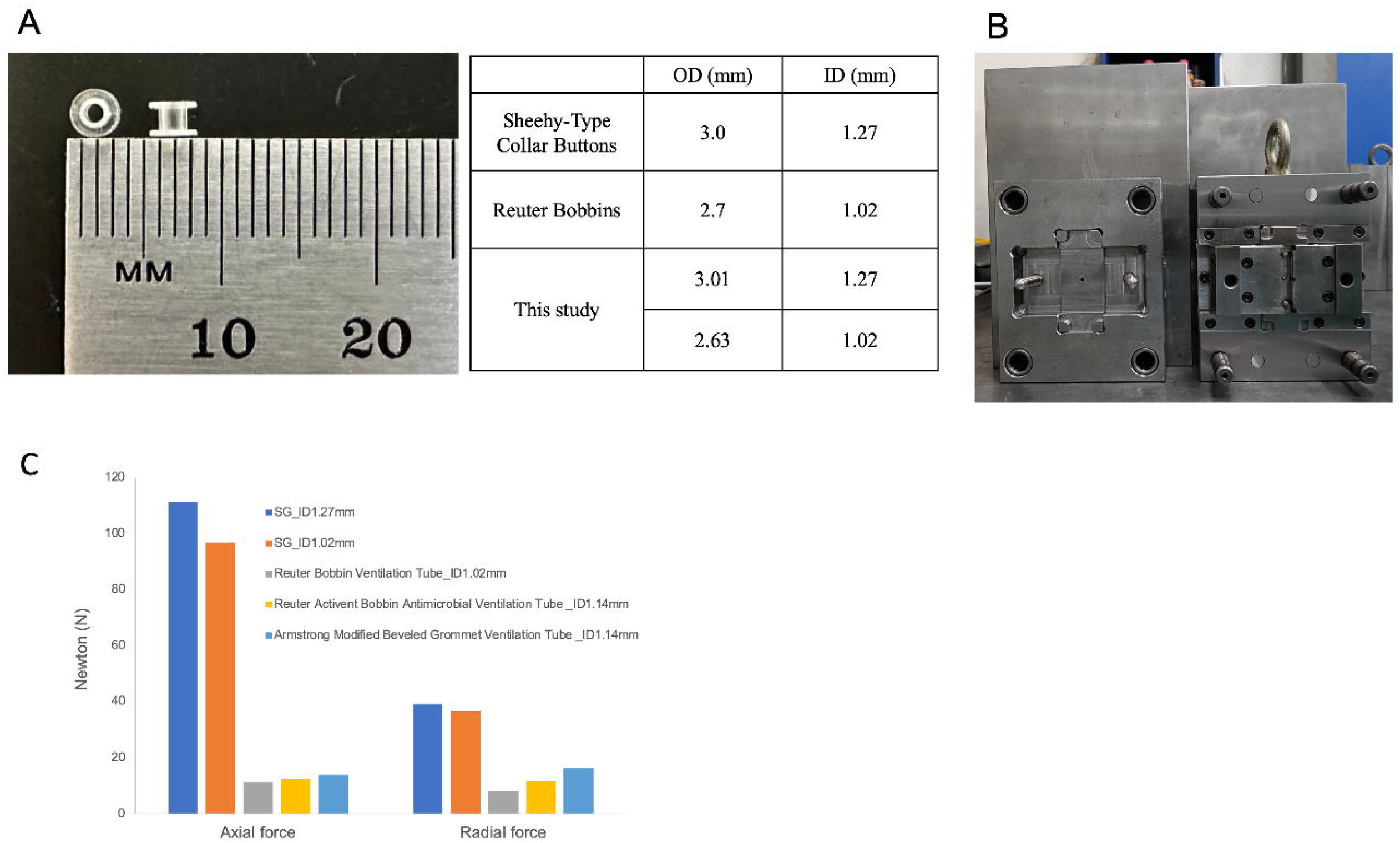
Design and mechanical properties of the bioabsorbable middle ear ventilation Tubes. (A) Image of the custom-designed PLA material alongside a ruler, indicating its dimensions. The table provides a comparison of the outer diameter (OD) and inner diameter (ID) of the PLA material used in this study (3.01 mm OD, 1.27 mm ID, and 2.63 mm OD, 1.02 mm ID) with standard Sheehy-Type Collar Buttons (3.0 mm OD, 1.27 mm ID) and Reuter Bobbins (2.7 mm OD, 1.02 mm ID). (B) The medical-grade stainless steel mold developed to enable the injection molding process. (C) Pressure test demonstrating superior strength of the PLA tube upon axial and radial forces compared to other products.

### 2.2 Mechanical and Biocompatibility Assessments

The product was packaged using Tyvek, and gamma irradiation was chosen as the sterilization method. A dose audit subsequently confirmed the sterility of the packaged products, with an applied dose ranging from 10 to 15 kGy. In terms of biocompatibility, the ventilation tube was categorized as Category C according to ISO 10993 standards. Both cytotoxicity and sensitization tests were completed and yielded acceptable results. The mechanical strength of the PLA tube was evaluated using a pressure test that applied axial and radial forces to the product. Overall, the product fulfills the sterility, mechanical, and biocompatibility requirements, from raw material selection through manufacturing, sterilization, and packaging, making it suitable for subsequent animal studies.

### 2.3 Animal Model Development and In Vivo Safety Evaluation

Guinea pigs were housed at the Laboratory Animal Center of the College of Medicine, National Taiwan University. All experimental procedures were conducted in accordance with institutional animal welfare guidelines and were approved by the Institutional Animal Care and Use Committee (IACUC) of the National Taiwan University College of Medicine (Approval No. 20240003).

A comprehensive *in vivo* model was established using guinea pigs weighing 200–250 g for the evaluation of middle ear implantation. Surgical techniques were developed for placing materials into the middle ear cavity. Anesthesia was induced using 3% inhalational isoflurane and maintained at a level sufficient to keep the animals sedated during the procedure and auditory brainstem response (ABR) testing. The surgical site was prepared by shaving the fur behind the left ear to expose the skin (Figure 2). The contralateral ear served as a surgical control (Mock), in which the mastoid portion of the ventral tympanic bulla was opened without implanting a ventilation tube. Under microscopic magnification, a 1–1.5 cm post-auricular skin incision was made using a scalpel. Blunt dissection was performed through the subcutaneous fat layer, which varied in thickness among animals. The bulla was accessed by gently rotating a scalpel to pierce the posterior-superior aspect of the ridge of the tympanic bulla. A small fenestration was created in the mastoid portion of the ventral bulla, and the ventilation tube was inserted into the middle ear cavity using forceps (Figure 2).

**Figure 2.**
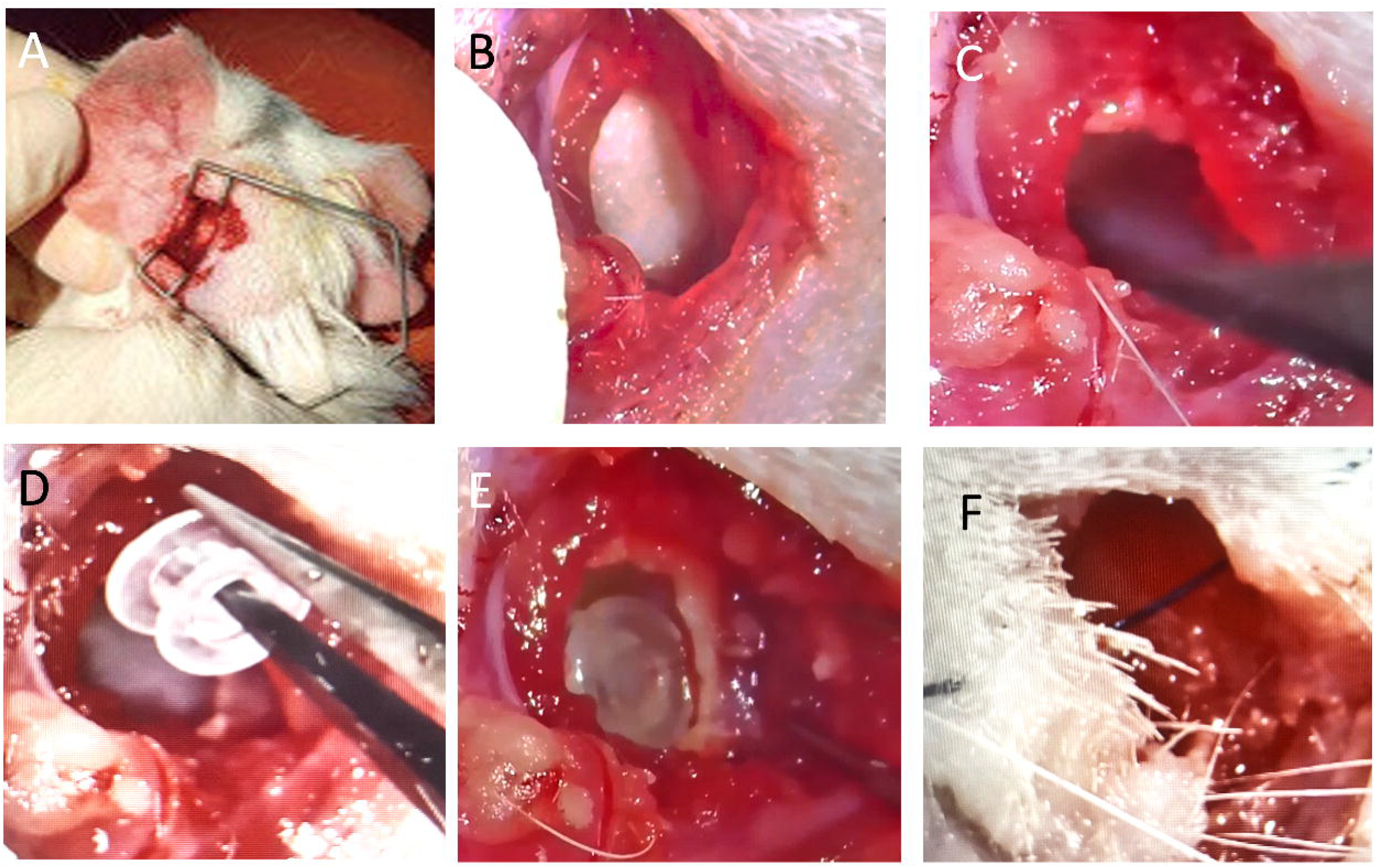
Surgical implantation of PLA tubes in the guinea pig model. (A) Preparation for surgery, with the ear secured. (B) A closer view of the surgical site, revealing the incision. (C) Use of a probe within the incision. (D) Insertion of the PLA material using forceps. (E) PLA tube in place within the surgical site. (F) The ear after the procedure, with the incision closed.

### 2.4 Audiological Evaluations

ABR thresholds were assessed in guinea pigs using an evoked potential detection system (Smart EP 3.90, Intelligent Hearing Systems, Miami, FL, USA), as previously described by Lu et al. (2011). Auditory stimuli, including click sounds and tone bursts at 8, 16, and 32 kHz, were presented at varying intensities from 10 to 90 dB SPL. ABR responses were recorded using subcutaneous needle electrodes inserted ventrolaterally near the ears. Waveforms and thresholds were measured and monitored throughout the observation period, with comparisons made to baseline levels to evaluate changes in auditory function.

### 2.5 Imaging and Histology Studies

Otoscopic examination was performed to assess the general condition of the tympanic membrane and external auditory canal prior to further imaging. Micro-computed tomography (micro-CT) imaging was employed to provide non-destructive, in vivo visualization of the implanted tube’s position, structural integrity, and any early signs of degradation (Park et al., 2011). Representative CT scans were obtained periodically, up to 9 months post-implantation. Gross anatomical evaluation was conducted during necropsy to assess macroscopic tissue responses and implant site morphology. For histopathological analysis, cochleae were harvested at 12 months postoperatively. Samples were fixed in 10% paraformaldehyde for 48 hours, decalcified, embedded in paraffin, sectioned into 8-μm slices, and stained with hematoxylin and eosin (H&E).

## 3. Results

### 3.1 PLA Middle Ear Ventilation Tube Design, Fabrication, Mechanical Properties, and Biocompatibility

Based on a detailed analysis of existing commercial products (e.g., Sheehy-type collar buttons, Medtronic), two bioabsorbable middle ear ventilation tube prototypes with different sizes were successfully designed and fabricated. Figure 1A presents the design specifications, including dimensions and structural characteristics. These designs informed the development of specialized molds (Figure 1B), which were essential for efficient production. Through the optimization of raw material cleaning, drying procedures, and injection molding parameters, high-quality PLA tubes were consistently produced. Importantly, pressure tests demonstrated the superior strength of the PLA tube when subjected to axial and radial forces compared to other products, as shown in Figure 1C.

The PLA middle ear ventilation tubes developed in this study have successfully passed preliminary evaluations in injection molding, packaging, sterilization, and biocompatibility (cytotoxicity and sensitization tests). This demonstrates its suitability not only as a test sample for the animal experiments in this study but also as a potential candidate for a formal commercialization process in the future.

### 3.2 Animal Model Development

A comprehensive animal model in guinea pigs was established for in vivo evaluation. Figure 2 illustrates the successful development of surgical techniques for implanting the larger PLA prototype (3.01 mm OD, 1.27 mm ID) into the middle ear cavity. Initial safety evaluations involved inserting the ventilation tube into the middle ear cavity by opening the mastoid portion of the ventral tympanic bulla.

### 3.3 Audiological Results

ABR testing was performed to evaluate the effects of PLA tube implantation on hearing thresholds over time. As shown in Figure 3, guinea pigs in the control, mock, and PLA groups (n = 5 per group) underwent audiological assessments at 1, 6, and 12 months post-operation. Both the mock and PLA groups maintained normal hearing thresholds throughout the 12-month period, indicating that neither the presence of the PLA tube in the middle ear nor the surgical procedure itself caused significant hearing loss or adversely affected auditory function. ABR waveforms and thresholds remained stable and comparable to baseline values, suggesting that PLA tube implantation preserved normal middle ear function, which is essential for efficient sound conduction during the entire observation period.

**Figure 3.**
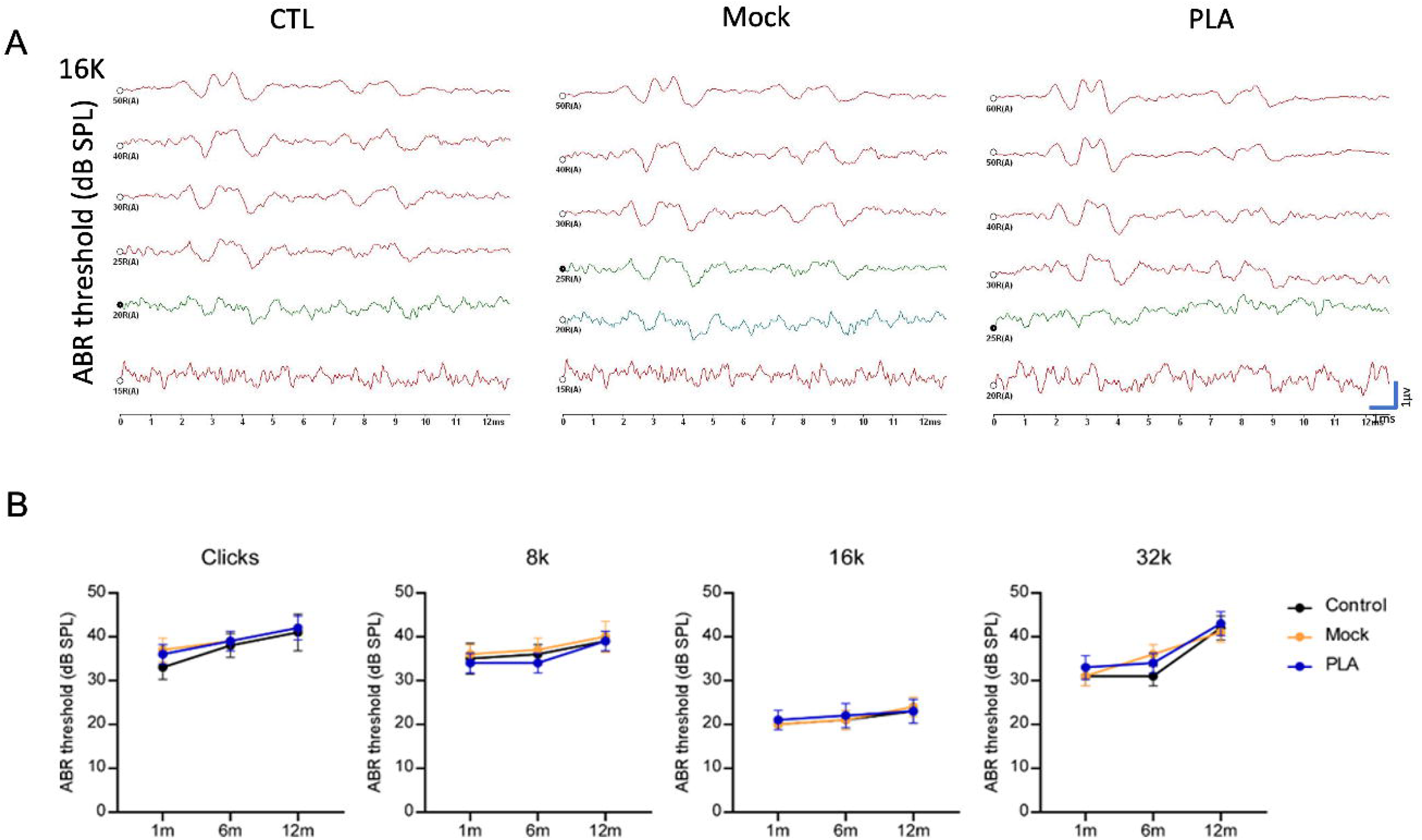
Longitudinal hearing results after PLA tube implantation. (A) Representative auditory brainstem response (ABR) waveforms at decreasing sound pressure levels (decibel sound pressure level, dB SPL) for the control, mock, and PLA groups, respectively, with 16K stimulus, showing positive waveforms at 20 dBPSL in all groups. (B) Comparison of ABR thresholds in dB SPL for the control, mock, and PLA groups over 1, 6, and 12 months for each frequency, demonstrating no difference between the three groups.

### 3.4 Imaging and Histology Results

Otoscopic examinations performed at 1, 6, and 12 months post-implantation revealed no visible inflammation or adverse reactions in the tympanic membrane or middle ear at any time point up to 12 months (Figure 4A). Specifically, no evidence of hemotympanum or tympanic membrane perforation was observed in any of the groups, and the tympanic membranes in the all groups remained normal throughout the study duration. These observations support the *in vivo* biocompatibility and mechanical stability of the PLA material.

**Figure 4.**
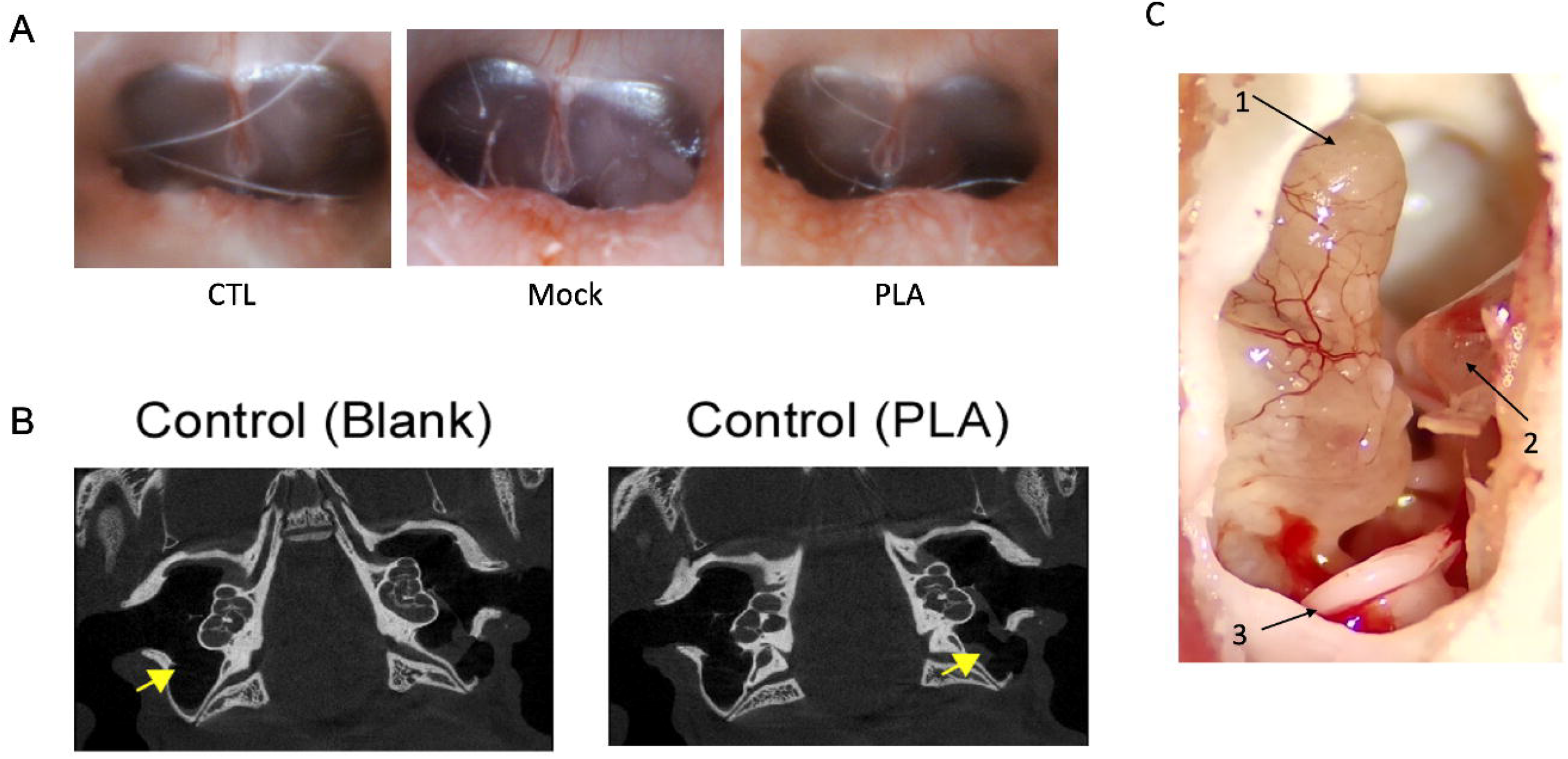
Otoscopic examination and micro CT scans. (A) Representative otoscopic images of the tympanic membrane at 12 months after implantation of the PLA tube, demonstrating their stable position and absence of visible inflammation or adverse reactions. (B) Micro-computed tomography (micro-CT) images taken up to 9 months post-implantation, illustrating the sustained position and structural integrity of the implanted PLA tubes within the middle ear cavity, along with early signs of material degradation. (C) Gross anatomical assessment at necropsy at 12 months demonstrating intact middle ear, intact tympanic membrane, and minimal foreign body response at the implant site. 1. cochlear apex 2. tympanic membrane 3. PLA

Non-invasive micro-computed tomography (micro-CT) imaging provided essential longitudinal insights into the tube’s position, structural integrity, and the onset of material degradation. Representative micro-CT images (Figure 4B), obtained up to 9 months post-implantation, confirmed the preservation of tube morphology within the middle ear cavity, along with early signs of gradual biodegradation over time. Gross anatomical assessment at necropsy further corroborated these findings, revealing intact middle ear, intact tympanic membrane, and minimal foreign body response at the implant site (Figure 4C).

## 4. Discussion

### 4.1 Summary of Key Findings

In this study, we successfully designed, fabricated, and preclinically evaluated a novel bioabsorbable middle ear ventilation tube utilizing PLA as the primary material. Our results demonstrate the successful development of the prototype with dimensions comparable to commercially available non-absorbable tubes, confirming the feasibility of the manufacturing process using optimized injection molding techniques. Importantly, *in vitro* mechanical and biocompatibility assessments, including pressure, cytotoxicity and sensitization tests, confirmed the material’s safety profile. The *in vivo* evaluation in a guinea pig model further validated the safety and biocompatibility of the PLA material, showing no significant inflammation or adverse reactions. Crucially, ABR measurements indicated that the implanted PLA tubes did not adversely affect hearing function. Advanced micro-CT imaging provided *in vivo* evidence of the tube’s sustained position and initial degradation over 9 months, while histological analyses corroborated the favorable tissue response and absence of significant foreign body reactions. These findings collectively establish our PLA middle ear ventilation tube as a promising solution with appropriate biocompatibility and an observable degradation profile.

### 4.2 Advantages over Existing Products

A key innovation of this study is the development of a bioabsorbable PLA middle ear ventilation tube, offering significant advantages over traditional non-absorbable devices (e.g., Teflon, silicone, stainless steel). Current devices often necessitate secondary removal surgery due to complications like extrusion failure, granulation tissue, tympanosclerosis, and persistent tympanic membrane perforation [14, 17]. In contrast, our PLA tube is designed to absorb within the middle ear over a clinically relevant timeframe (1-2 years). This aims to eliminate the need for additional interventions, reducing patient risks, healthcare costs, and long-term tympanic membrane complications.

Several types of biomaterials have been applied to develop absorbable middle ear ventilation tubes (Table 1). D’Eredità and Marsh (2005) found that poly-bis(ethylalanate)phosphazene tubes in guinea pigs showed excellent tympanic membrane healing with no infection or inflammatory reaction, and their disintegration rate could be controlled by varying the polymer formulation [13]. Massey et al. observed in a guinea pig model that PLA tubes remained in the tympanic membrane longer (average 63.2 vs. 19.3 days) than 50/50 PLA/PLGA tubes (average 18.8 vs. 8.1 days), with both demonstrating good biocompatibility and complete tympanic membrane healing [18]. Ludwick et al. showed that PLA tubes possessed bacteriostatic properties against *Pseudomonas aeruginosa* and *Staphylococcus aureus* in vitro [17], suggesting a potential reduction in post-placement otorrhea. Park et al. further explored polyester tubes in chinchillas, noting that while PLGA tubes showed more inflammation, silver-coated PLGA tubes had significantly less fibrosis and lower neutrophil counts [14], indicating an anti-inflammatory effect. Skovlund et al. conducted a human feasibility study, demonstrating that a novel bioabsorbable gelatin ventilation tube provided intermediate-term middle ear ventilation for an average of 12 weeks, supporting its clinical applicability [19].

**Table 1.**
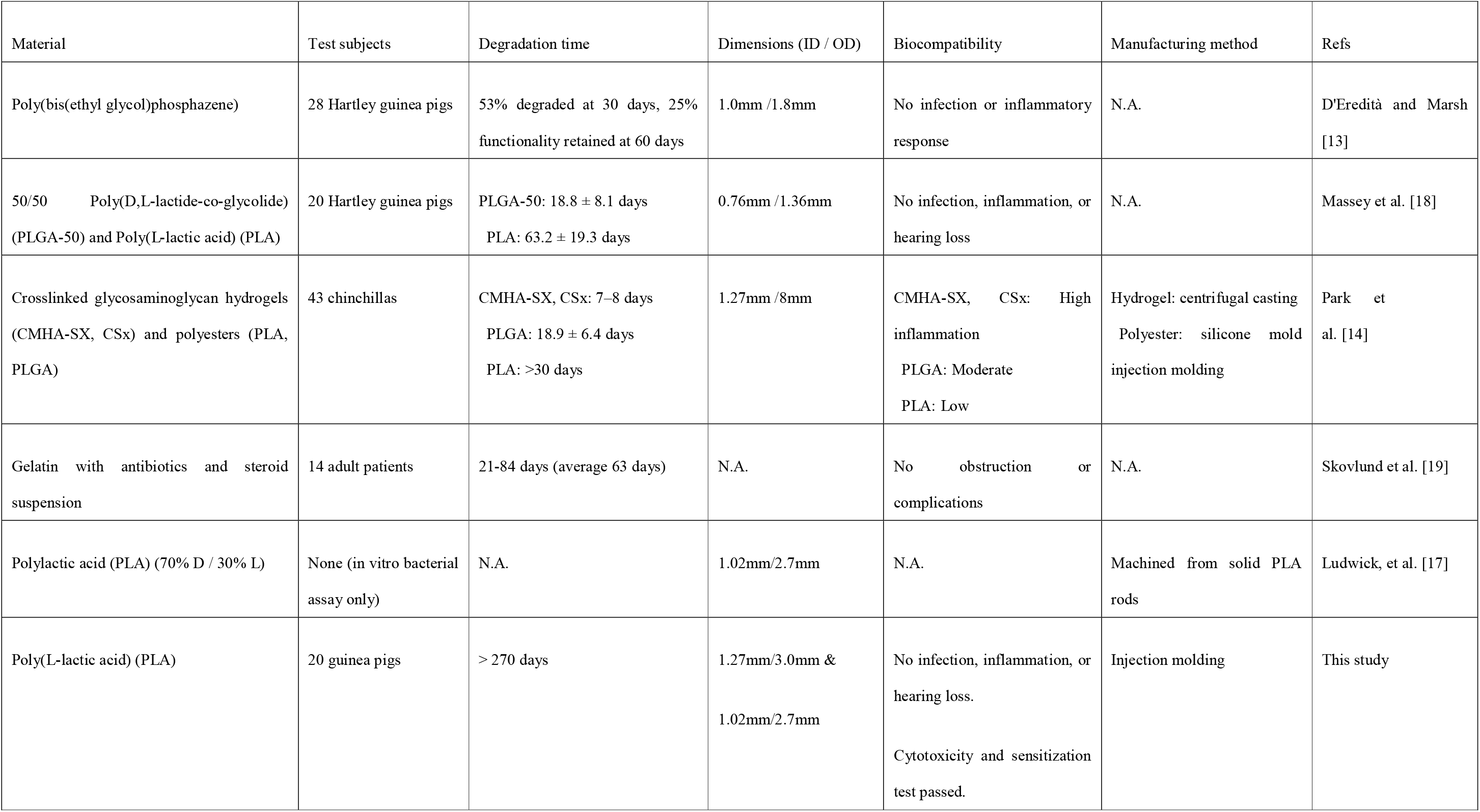
Comparison of existing absorbable middle ear ventilation tube products.

Compared to previous studies, our research on a novel bioabsorbable PLA middle ear ventilation tube in guinea pigs demonstrates notable strengths. Earlier poly-bis(ethylalanate)phosphazene tubes degraded within 30 to 60 days, PLA/PLGA tubes were resorbed within 18.8 to 63.2 days in guinea pigs and within 18.9 days for PLGA and over 30 days for PLA in chinchillas, but our PLA tubes exhibited sustained structural integrity, with initial signs of degradation appearing after nine months (over 270 days). This prolonged, controlled degradation profile *in vivo* is specifically designed to match the ideal 1–2-year clinical ventilation period and represents a considerable advantage over prior materials that often degrade too rapidly. Furthermore, our PLA tubes demonstrated excellent biocompatibility. There was no significant inflammation, foreign body reaction, or negative impact on hearing function, as measured by ABR. This is consistent with previous PLA findings. Unlike some cross-linked hydrogel tubes, which caused high inflammation and enlarged the myringotomy site, our PLA design avoids such adverse responses. Successful fabrication via injection molding ensures consistent quality and suitability for mass production. This comprehensive preclinical evaluation showing favorable biocompatibility and maintained hearing, as well as a significantly extended and controlled degradation time, positions our PLA ventilation tube as a highly promising alternative that could eliminate the need for secondary removal surgeries and reduce associated complications in human patients.

### 4.3 Correlation with In Vitro and In Vivo Data

The robust *in vitro* material characterization and processing optimization were instrumental in guiding the development of PLA prototypes with favorable *in vivo* performance. Stringent controls during raw material handling and injection molding ensured consistent tube production with desired properties and confirmed the superior mechanical strength of the PLA tube. While long-term *in vitro* degradation and mechanical strength data are ongoing, preliminary *in vivo* micro-CT imaging showed sustained presence and initial signs of absorption over 9 months, aligning with our 1–2-year target. Maintained hearing function and minimal inflammatory response histologically underscore the successful translation of in vitro biocompatibility to the complex in vivo middle ear environment.

### 4.4 Future Directions and Clinical Translation

While the preclinical data is promising, further *in vitro* degradation and mechanical strength tests are essential to fully understand the long-term performance and absorption kinetics of the PLA tubes. Furthermore, future research will explore advanced functionalities, such as antibacterial coatings applied via dip-coating methods, to develop an “absorbable antibacterial middle ear ventilation tube.” This enhancement could further mitigate the risk of postoperative infections. The next step involves initiating human clinical trials to evaluate the safety and efficacy of the PLA tubes, with the ultimate goal of improving patient outcomes and reducing the complications associated with current non-absorbable devices.

## 5. Conclusion

This study successfully developed and preclinically evaluated a novel bioabsorbable PLA middle ear ventilation tube. The prototype demonstrated comparable dimensions to existing commercial tubes, confirmed *in vitro* biocompatibility, and was well-tolerated in a guinea pig model without adverse reactions or impact on hearing. Micro-CT imaging and histological analyses showed structural integrity and initial degradation over 9 months, supporting the targeted degradation profile. These findings indicate the PLA tube is a promising alternative to non-absorbable tubes, potentially eliminating secondary surgeries and reducing complications. Preclinical data support further clinical trials to validate its safety and efficacy in human patients.

